# Age-related declines in neural distinctiveness correlate across brain areas and result from both decreased reliability and increased confusability

**DOI:** 10.1101/2021.06.28.449009

**Authors:** M. Simmonite, T. A. Polk

## Abstract

According to the neural dedifferentiation hypothesis, age-related reductions in the distinctiveness of neural representations contribute to sensory, cognitive, and motor declines associated with aging: neural activity associated with different stimulus categories becomes more confusable with age and behavioural performance suffers as a result. Initial studies investigated age-related dedifferentiation in the visual cortex, but subsequent research has revealed declines in other brain regions, suggesting that dedifferentiation may be a general feature of the aging brain. In the present study, we used functional magnetic resonance imaging to investigate age-related dedifferentiation in the visual, auditory, and motor cortices. Participants were 58 young adults and 79 older adults. The similarity of activation patterns across different blocks of the same condition was calculated (within-condition correlation, a measure of reliability) as was the similarity of activation patterns elicited by different conditions (between-category correlations, a measure of confusability). Neural distinctiveness was defined as the difference between the mean within- and between-condition similarity. We found age-related reductions in neural distinctiveness in the visual, auditory, and motor cortices, which were driven by both decreases in within-category similarity and increases in between-category similarity. There were significant positive cross-region correlations between neural distinctiveness in different regions. These correlations were driven by within-category similarities, a finding that indicates that declines in the reliability of neural activity appear to occur in tandem across the brain. These findings suggest that the changes in neural distinctiveness that occur in healthy aging result from changes in both the reliability and confusability of patterns of neural activity

## Introduction

Aging, even in the absence of disease, is associated with declines in sensory, cognitive, and motor function. Influential computational models of aging attribute some of these changes to a phenomenon known as age-related neural dedifferentiation (Li cites), in which neural representations of different stimuli become less distinctive with age. The earliest empirical support for neural dedifferentiation came from positron emission tomography studies that reported age-related reductions in the functional specialization of the ventral and dorsal visual pathways (Grady et al., 1992, 1994). Using fMRI, Park et al., (2004) found that ventral visual activity patterns in response to faces, houses, and words were more similar to each other (less distinctive) in older vs. younger adults. For example, older adults exhibited only slightly greater activation in the fusiform face area (FFA) when viewing faces, compared to when viewing words or houses, while young adults exhibited much more selective activation.

Subsequent studies exploited the ability of multivariate pattern analysis (MVPA) approaches to identify fine-grained differences in patterns of neural activity and added to the evidence for age-related declines in neural distinctiveness. Carp et al., (2011) evaluated the distinctiveness of distributed patterns of neural activation elicited by different categories of visual stimuli and found neural activation patterns in the ventral visual cortex were less distinct in older adults. Several other studies have used MVPA to replicate these findings of reduced age-related dedifferentiation at the level of category representations (Chamberlain et al., 2021; J. Park, Carp, Hebrank, Park, & Polk, 2010) and recently Kobelt et al., (2021) found that item-level distinctiveness was also reduced in older adults.

Many of the early studies focused on neural representations of visual categories and demonstrated age-related neural distinctiveness reductions in areas of the visual cortex, but other research has indicated that a similar effect can be observed in other sensory regions. For example, recent studies have shown age-related reductions in neural distinctiveness in the auditory (Du, Buchsbaum, Grady, & Alain, 2016; Lalwani et al., 2019) and motor (Carp, Park, Hebrank, Park, & Polk, 2011; Cassady, Gagnon, et al., 2020) cortices.

There is also growing evidence supporting the role of age-related declines in neural distinctness during higher order cognitive processes, such as memory. Carp et al., (2010) found evidence for such declines in the prefrontal and parietal cortices during maintenance of high memory loads. Links have also been discovered between age-related changes in neural representation and episodic memory performance (Koen, Hauck, & Rugg, 2019; Zheng et al., 2018), working memory encoding (Payer et al., 2006) and recognition performance (Berron et al., 2018).

Recent work has also begun to explore the mechanisms underlying age-related neural dedifferentiation. For example, several studies have investigated whether dedifferentiation is the result of neural broadening (i.e., brain regions that are relatively category-selective in younger adults respond more to non-preferred stimuli in older adults), neural attenuation (i.e., category-selective brain regions that respond strongly to preferred stimuli in young adults respond less strongly in older adults), or both. Evidence for all three possibilities has been observed. Koen et al., (2019) found evidence that neural attenuation drove their findings of reduced neural distinctiveness in older adults. On the other hand, Kobelt et al., (2021) found increased neural activation in response to non-preferred stimuli in older adults, with no age differences in activation to preferred stimuli, supporting the neural broadening hypothesis. Park et al., (2012) provided support for both, finding evidence of neural broadening in the FFA and neural attenuation in the extended face network. Evaluating this evidence, Koen and Rugg (2019) conclude that different mechanisms underlie age-related neural dedifferentiation in different brain regions.

In this study, we explore another question about mechanism: Is neural dedifferentiation due to age-related declines in the reliability of neural activation patterns (reduced within-category similarity), to increased confusability of activity in response to different stimulus categories (increased between-category similarity), or both. Carp et al., (2011) found age differences in both, with older adults exhibiting reduced within-category similarities and increased between-category similarities. Other evidence has demonstrated age-reductions in the reliability of representations at the level of individual items (Zheng et al., 2018) and categories, without a significant increase in between-category similarity (Chamberlain et al., 2021). Here we use data from the Michigan Neural Distinctiveness (MiND) project (Gagnon et al., 2019) to investigate that question in visual cortex, in auditory cortex, and in motor cortex.

We also use this dataset to explore another question related to mechanism that has not previously been investigated: Is dedifferentiation in one brain region associated with dedifferentiation in other regions. For example, do older adults with less distinctive visual representations also exhibit less distinctive motor and auditory representations? This issue has important implications for theoretical models of cognitive aging: while common-cause theories argue that age-related declines occur in tandem across domains, process-specific theories predict instead that different abilities decline independently (D. C. Park et al., 2002).

## Materials and Methods

### Participants

Participants were 58 young adults (mean age 22.76 ± 2.86, range 18 – 29 years; 29 females) and 79 older adults (mean age 70.44 ± 5.04, range 65 – 87; 52 females) who were recruited to take part in the study from the Ann Arbor community. Data were collected as part of the Michigan Neural Distinctiveness (MiND) project, a large multi-modal research project which is described in Gagnon et al., (2019).

All participants were healthy, right-handed, native English speakers with normal hearing and normal or corrected-to-normal vision. Prior to enrollment in the study, participants took part in a telephone screening to ensure that they were free from MRI safety contraindications and were not taking medications with vascular or psychotropic effects. Additionally, all participants were free of significant cognitive impairment, with an overall cognition score of 85 or greater as measured using the NIH Toolbox Cognition Battery (Weintraub et al., 2013).

All participants gave full written informed consent prior to their participation in the study. The study protocol was approved by the Institutional Review Board of the University of Michigan.

### Study Design

Participants enrolled in the study completed three separate testing sessions: a behavioral testing session, a functional magnetic resonance imaging (fMRI) session, and a magnetic resonance spectroscopy (MRS) session. Participants completed these sessions on different days and completed all three within an average of 22 days. Here we describe the fMRI protocol, other details and parameters of the MiND study are provided in Gagnon et al., (2019).

### Functional MRI Tasks

While fMRI data were being collected, each participant performed one run of each of three tasks, each designed to elicit neural activity in a different brain region – visual, auditory, and motor cortex. Each of the three tasks lasted 6 minutes, had two experimental and one control condition, and followed the same block design format – six 20 sec blocks of each of the experimental conditions, and twelve 10 sec control blocks with no stimulus. Each experimental block was followed by a control block and task blocks were pseudorandomized, with block order the same for all participants.

The visual task was based on previous studies of age-related dedifferentiation (Carp, Park, Polk, et al., 2011; D. C. Park et al., 2004; J. Park et al., 2012) and consisted of experimental blocks in which photographs of male faces or houses were presented for 20 seconds. Experimental blocks contained 20 stimuli each presented for 500 ms with a 500ms interstimulus interval during which a fixation cross was presented. Control blocks contained a fixation cross presented for 10 sec. To ensure participants were paying attention during the viewing of the images, they were instructed to press a button with their right index finger when they saw a target. During face blocks, targets were images of women’s faces, and during house blocks, targets were images of apartment buildings.

The auditory task consisted of experimental blocks in which foreign speech or instrumental music was presented for 20 seconds. Each speech block contained a segment of a news reporter speaking in a different language (Creole, Macedonian, Marathi, Persian, Ukrainian and Swahili). During screening it was ensured that participants were not familiar with any of these languages. Control blocks contained no sound. A fixation cross was presented onscreen for the duration of the task. Participants were asked to respond to targets (a beep presented alongside the auditory stimuli) by pressing a button with their right index finger.

The motor task consisted of experimental blocks in which left-pointing or right-pointing arrows were presented. Experimental blocks consisted of one type of arrow – left or right – only. Within each block, 20 arrows were presented each for 500 ms with a 500 ms ISI during which a fixation cross was presented. Control blocks lasted for 10 seconds and consisted of a fixation cross. To ensure participants were attending to the sounds, they were asked to make a button press with their left thumb each time they saw the left-pointing arrow stimulus and with their right thumb each time they saw the right-pointing arrow. Since the motor task required active responses, this task did not include targets.

In the visual and auditory tasks, targets were presented approximately once per minute, and there was never more than one target in each block. Responses were collected using a Celeritas 5-button fiber-optic response unit and sound was presented through an Avotec Conformal Headset.

### MRI acquisition

Structural and functional brain images were acquired at the University of Michigan’s Functional Magnetic Resonance Imaging Laboratory, using a GE Discovery MR750 3T MRI scanner with a GE 8-channel head coil. Participant motion was minimized using cushions and Velcro straps. A high-resolution anatomical image was obtained using a 3D fast spoiled gradient-echo acquisition (SPGR) BRAVO sequence with the following parameters: repetition time (TR) = 12.2 ms; echo time (TE) = 5.2 ms; inversion time (TI) = 500ms, flip angle = 15°; field of view = 256 × 256 and voxel size 1 × 1 × 1 mm (156 axial slices). Functional images were acquired using a single shot gradient-echo reverse spiral pulse sequence with the following parameters: TR = 2000ms; TE = 30ms, flip angle = 90°, field of view = 220 × 220 mm; 180 volumes and voxel size 3 × 3 × 3 mm (43 axial slices). The duration of each functional scan run (i.e., each task) was 6 minutes.

### MRI preprocessing

Functional data were k-space despiked, reconstructed and corrected for physiological motion effects (respiration and cardiac-induced noise) using RETROICOR. Data were slice-time corrected using SPM8 and motion corrected using FSL’s mcflirt. All further analysis steps were performed using Freesurfer. Data were resampled into surface space, based on a white/grey matter segmentation of the participants high resolution anatomical image, and smoothed using a 5mm 2D smoothing kernel.

### ROI definition

For each of the three tasks, individualized ROI masks for each participant were created in two steps. First, each participant’s structural image was segmented using FreeSurfer’s Cortical Parcellation tool, and bilateral anatomical masks were created by combining cortical regions that were hypothesized to be relevant for that task. For the visual task, this consisted of the bilateral fusiform gyrus and bilateral parahippocampal gyrus; for the auditory task, this consisted of the bilateral superior temporal gyrus, the bank of the superior temporal sulcus, the transverse temporal gyrus and the supramarginal gyrus; and for the motor task this consisted of the precentral gyrus, postcentral gyrus and supramarginal gyrus.

Second, we estimated neural responses by fitting a GLM for each task, implemented in Freesurfer’s FSFAST pipeline. Each model included regressors for the two experimental conditions (visual task: faces and houses; auditory task: speech and instrumental music; motor task: left and right thumb button presses) and the control condition, each of which were convolved with a standard hemodynamic response function. Contrasts were then specified for each of the experimental conditions relative to the control condition. These contrasts were then used to construct functional ROIs for each participant.

For example, for the visual task, we took the face vs. control and house vs. control contrast, selected the vertices that fell within that individual’s anatomical ROI and sorted the beta values at these vertices in descending order. Using these two lists of sorted beta values, we alternated adding the most active vertex from each list (i.e., the highest beta value) to the ROI, followed by the next most active vertex, and so on. If a vertex had already been added to the ROI because of high activation in the other condition, we then added the next most active vertex from that condition to the ROI. This elicited functional ROIs that were comprised of the most active vertices during the task and included an equal number of vertices from the two experimental conditions.

We used this method to produce ten ROIs of varying size and constructed them so that there was no overlap between the ROI’s (i.e., the first ROI consisted of the 50 most active vertices, the second included 51 – 100, and then 100-200, until we had 10 ROIs). This meant that each ROI was independent of all others, and no one vertex was included in more than one ROI. An additional, large ROI containing the 2000 most active vertices within the anatomical mask was also created, which was used to calculate correlations between distinctiveness measures in different brain regions.

A similar process was performed for each of the fMRI tasks, resulting in person-specific ROIs for the visual task, the auditory task, and the motor task.

### Distinctiveness calculation

To assess neural distinctiveness, we used a correlation-based approach, in line with that used by Haxby et al. (2000), Carp et al. (2011), Lalwani et al. (2019), Cassady et al. (2020), and Chamberlain et al. (2021). To do so, for each participant, in each task we compared patterns of neural activity that were elicited by different blocks of the same experimental condition, with neural activity elicited by blocks of different experimental conditions.

For each participant, we estimated neural responses for each task by fitting a GLM. This model was separate to the one outlined in the section above, which was used for ROI creation. In the current GLM, separate regressors were included for each of the 12 experimental blocks included in the task, resulting in 12 beta values at each vertex – each one estimating activity during one experimental block.

Using the beta values for vertices within a functional ROI, we calculated correlations between the beta values of all pairs of experimental blocks of the same type, then calculated the mean of these correlations to get a value which describes the average within-category similarity for that task. We also calculated correlations between the beta values of all unique combinations of pairs of experimental blocks of different categories (within the same task) and averaged these values to get a value that describes the average between-category similarity for that task. We then subtracted the between-category similarity from the within-category similarity to obtain a measure of neural distinctiveness. This value has a theoretical range of -2 (indicating that the neural representations of that participant are more similar between different categories than they are within the same category, therefore demonstrating low neural distinctiveness) to +2 (indicting that the neural patterns of that participant show a higher degree of similarity when elicited by the same category than they do for different categories, therefore demonstrating high neural distinctiveness). This was repeated for all 10 ROI sizes and each of the tasks.

### Statistical analysis

All statistical analysis was performed in R. For all measures, outliers greater than three standard deviations above or below the age group mean were removed. To investigate group differences in neural distinctiveness across the different ROIs, we used multilevel models with age group as a between-subjects factor (two levels: young and older) and ROI size as a within-subjects factor (ten levels, ranging from the 50 most activated vertices to the 5000-10000 most activated vertices). Separate models were performed for each of the three tasks (visual, auditory, and motor), for each of the three measurement types (distinctiveness, within and between).

To investigate the relationship between distinctiveness measures in different brain regions, we calculated partial correlations using the 2000 vertex ROIs. Relationships were also investigated separately in the young and older age groups using bivariate correlations.

## Results

Technical difficulties with auditory equipment meant that fMRI data during the auditory task were not acquired for two older participants. Auditory fMRI data from one young and one old participant were also excluded as they performed a previous version of the task. We excluded any fMRI data during which the participant moved more than 2mm translation or 2 degrees rotation. This led to the exclusion of three young participants’ motor data, two young participants’ visual data, one older participants’ auditory data and six older participants’ visual data.

### Age differences in neural distinctiveness

Neural distinctiveness measures for all tasks are presented in Figure 1, and full results of multilevel models are included in Table 1. Consistent with previous findings, we found that the distinctiveness of neural representations was reduced in older adults when compared with younger adults, across all three tasks (visual: *x*^2^(1) = 13.29, *p*<.001; auditory: *x*^2^(1) = 21.14, *p*<.0001; motor: *x*^2^(1) = 5.11, *p*<.05). Additionally, there was a significant age × ROI size interaction effect for the measure of auditory distinctiveness (*x*^2^(9) = 31.87, *p* <.001) with larger ROIs exhibiting larger age effects.

**Figure 1:**
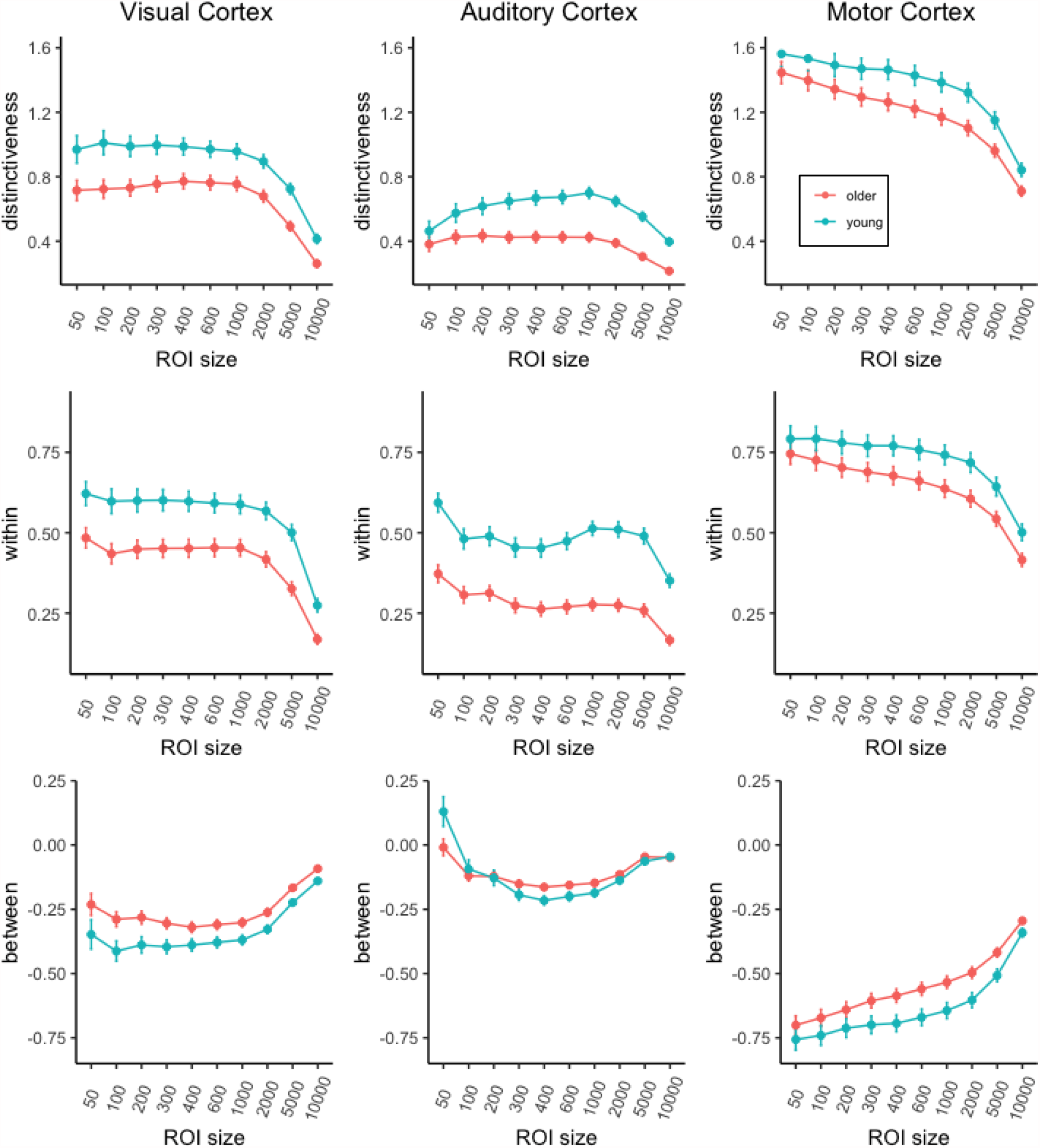
Age effects on neural distinctiveness measures as a function of ROI size in visual, auditory, and motor cortex. The top row plots neural distinctiveness, the middle row plots within-category similarity, and the bottom-row plots between-category similar

**Table 1:** Results from multilevel models investigating neural distinctiveness across age and ROI size

|  |  |  |  |  |  |  |  |  |
| --- | --- | --- | --- | --- | --- | --- | --- | --- |
|  |  | Main effect: Age |  |  | Main effect: ROI |  | Interaction effect:<br>Age x ROI |  |
| | | r | $\chi^2$ | p | $\chi^2$ | p | $\chi^2$ | p |
| Distinctiveness | Visual | .31 | 13.29 | .0003 | 535.46 | <.0001 | 7.53 | .58 |
|  | Auditory | .38 | 21.14 | <.0001 | 201.45 | <.0001 | 31.87 | .0002 |
|  | Motor | .19 | 5.11 | .02 | 1025.50 | <.0001 | 11.79 | .23 |
| Within | Visual | .33 | 15.08 | <.0001 | 683.85 | <.0001 | 8.04 | .53 |
|  | Auditory | .52 | 42.10 | <.0001 | 272.10 | <.0001 | 16.51 | .07 |
|  | Motor | .16 | 3.28 | .07 | 902.84 | <.0001 | 10.34 | .32 |
| Between | Visual | .26 | 8.72 | .0031 | 301.15 | <.0001 | 9.41 | .40 |
|  | Auditory | .02 | .07 | .79 | 312.35 | <.0001 | 51.46 | <.0001 |
|  | Motor | .19 | 5.13 | .02 | 1025.49 | <.0001 | 14.58 | .10 |

To determine whether reduced distinctiveness in older participants was driven by reduced similarity of neural activation patterns **within** blocks of the same stimulus category or increased similarity of neural patterns **between** categories, we repeated our analyses focusing on the within and between measures. Within category similarity measures were significantly reduced in older adults in the visual (*x*^2^(1) = 15.08, *p*<.0001) and auditory tasks (*x*^2^(1) = 42.10, *p*<.0001), are were marginally reduced in the motor task (*x*^2^(1) = 3.28, *p*=.07).

Between category similarities were significantly more negative in younger participants for the visual (*x*^2^(1) = 8.72, *p*<.01) and motor tasks (*x*^2^(1) = 5.13, *p*<.05), but not the auditory task (*x*^2^(1) = .07, *p* = .79). There was, however, a significant age × ROI size interaction effect for the measure of between category similarity in the auditory cortex (*x*^2^(9) = 51.46, *p* <.0001), an effect driven by a significant age difference in the 1-50 vertices ROI (t_90.30_ = 2.11, *p*<.05).

### Relationships between neural distinctiveness in different regions

Correlation coefficients between distinctiveness measures in the three regions are presented in Table 2 and Figure 2. Across all participants, controlling for age, there was a significant relationship between visual cortex distinctiveness and auditory cortex distinctiveness (*r* = .21, *p* <.05), and between visual cortex distinctiveness and motor cortex distinctiveness (*r* = .46, *p* <.001). When these relationships were investigated in the two age groups separately, the same pattern of significant relationships was found in the older participants (visual – auditory *r* = .24, *p* <.05; visual – motor *r* = .45, p <.001), however only the relationship between visual cortex distinctiveness and motor cortex distinctiveness was significant in the young (*r* = .49, *p* <.001).

**Table 2:** Correlation coefficients and sample sizes of cross-region distinctiveness relationships

|  |  | All participants:<br>partial correlations |  |  | Young participants:<br>univariate correlations |  |  | Older participants:<br>univariate correlations |  |  |
| --- | --- | --- | --- | --- | --- | --- | --- | --- | --- | --- |
|  |  | visual | auditory | motor | visual | auditory | motor | visual | auditory | motor |
| Distinctiveness | Visual |  | .21* | .46*** |  | .15 | .49*** |  | .24* | .45*** |
|  | Auditory | 124 |  | .09 | 55 |  | -.06 | 69 |  | .20<sup>†</sup> |
|  | Motor | 127 | 129 |  | 54 | 54 |  | 73 | 75 |  |
| Within | Visual |  | .51*** | .59*** |  | .42** | .68*** |  | .56*** | .54*** |
|  | Auditory | 124 |  | .43*** | 55 |  | .31* | 69 |  | .51*** |
|  | Motor | 127 | 129 |  | 54 | 54 |  | 73 | 75 |  |
| Between | Visual |  | -.02 | .18* |  | -.03 | .16 |  | .00 | .19 |
|  | Auditory | 124 |  | -.18* | 55 |  | -.19 | 69 |  | -.19 |
|  | Motor | 127 | 129 |  | 54 | 54 |  | 73 | 75 |  |

<sup>†</sup> $p < .10$ , \* $p < .05$ , \*\* $p < .01$ , \*\*\* $p < .001$
Note: correlation coefficients are presented above diagonal, $n$ is presented below the diagonal

**Figure 2:**
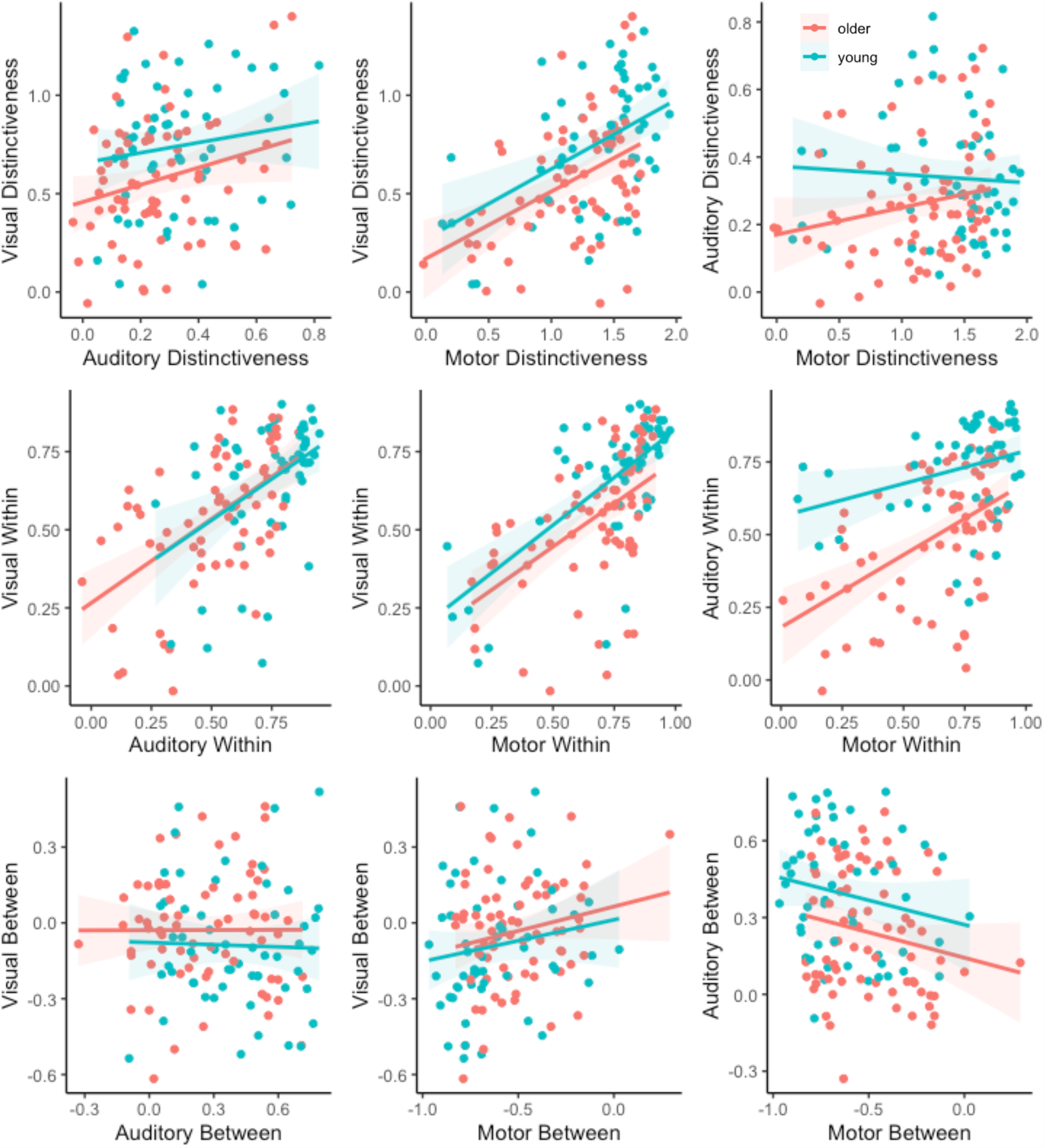
Scatterplots showing cross-region relationships in distinctiveness measures. The top-row plots neural distinctiveness, the middle-row plots within-category similarity and the bottom-row plots between-category similarities. Young adults are plotted in blue and older adults are plotted in red.

When we looked at cross-region relationships in within category similarities, there were significant correlations between the measures elicited by all three tasks (visual – auditory *r* = .51; visual – motor *r* = .59; auditory – motor *r* = .43, all *p*s <.001). These relationships were also significant in the young participant group (visual – auditory *r* = .42; visual – motor *r* = .68; auditory – motor *r* = .31, all *p*s <.05) and the old participant group (visual – auditory *r* = .56; visual – motor *r* = .54; auditory – motor *r* = .51, all *p*s <.001).

Finally, we looked at cross-region relationships in between category similarities. Across the complete participant sample, the only significant positive relationship across between category similarities was between the motor region and the visual region (*r* = .18, *p* <.05). There was also a significant relationship between the motor region and the auditory region, but it was negative (*r* = -.18, *p* < .05). Neither of these relationships were significant in the young or older subgroups when analyzed separately.

## Discussion

In this study, we investigated neural distinctiveness in the visual, auditory, and motor cortices of healthy young and older adults. In line with other reports, we found age-related declines in all three. When probing the mechanisms behind these reductions, we found that both the within-category similarity and between-category dissimilarity of neural representations was reduced in older vs. younger adults. Additionally, we found high cross-region correlations in neural distinctiveness in both young and older adults, which were driven by correlations in within-category similarity. We discuss each of these findings in turn.

### Age-related declines in neural distinctiveness result from both decreased within-category similarity and increased between-category similarity

Previous neuroimaging research has employed univariate and multivariate techniques to demonstrate age-related neural dedifferentiation in sensory regions including the visual (Carp, Park, Polk, et al., 2011; Chamberlain et al., 2021; D. C. Park et al., 2004; J. Park et al., 2012), auditory (Du et al., 2016; Lalwani et al., 2019), and motor (Carp, Park, Hebrank, et al., 2011; Cassady, Gagnon, et al., 2020) cortices, as well as in areas including the hippocampus (Yassa, Mattfeld, Stark, & Stark, 2011) inferior prefrontal cortex (Du et al., 2016) and perirhinal cortex (Berron et al., 2018). Consistent with these findings, we also found reduced neural distinctiveness in the visual, auditory, and motor cortices of older adults.

We also found significant age effects on within-category similarity in the visual and auditory cortices (and a marginally significant effect in the motor cortex), within-category similarity being lower in the older vs. younger adults. Put simply, young adults tended to produce similar activation patterns when the same stimulus category was presented repeatedly while older adults produced less similar patterns. One way of interpreting this finding is that activation patterns are less reliable in older adults, perhaps due to a noisier neural system. This interpretation is in line with prior discoveries of reductions in the fidelity of neural representations in older adults (Goh, Suzuki, & Park, 2010; St-Laurent, Abdi, Bondad, & Buchsbaum, 2014; Zheng et al., 2018).

We also found significant age-related declines in between-category dissimilarity in the visual and motor cortex (i.e., an increase in between-category similarity). That is, while faces and houses (and left and right button presses) elicited fairly different activation patterns in young adults, those same stimulus categories elicited more similar activation patterns in older adults. One interpretation is that neural representations are more confusable in older compared with younger adults, which could potentially undermine behavioral performance. Of course, less reliable within-category activation could also be associated with worse performance, so future studies could compare the extent to which within-category similarity and between-category similarity is associated with behavior in older adults.

Interestingly, despite these age-related increases in between-category similarity in the visual and motor cortex, we found no evidence for such an increase in auditory cortex. One possible explanation is differences in baseline between-category similarity in the young. The average between-category similarity of face and house activations in the young was significantly negative (average across all 10 ROIs = -0.34). Likewise, the average between-category similarity of left and right tapping activations in the young was also significantly negative (average across all 10 ROIs = -0.61). In contrast, the average between-category similarity of speech and music activations in the young was much closer to 0 (average across all 10 ROIs = -0.11. There was therefore much less room for the between-category similarity of the auditory activations to get more positive in the older adults relative to the visual and motor tasks. This interpretation would predict that age-related increases in between-category similarity would be observed in auditory cortex if auditory conditions were used that elicited more dissimilar activations in the young.

### Cross-region relationships in neural distinctiveness are driven by within-category similarities

When we investigated cross-region correlations in neural distinctiveness measures, we found a significant relationship between visual and motor cortex distinctiveness, which was significant both in young and older adults. When we explored cross-region relationships between within- and between-category correlations separately, it was apparent that the observed correlation between visual and motor cortex distinctiveness was driven by the strength of within-category similarities. Indeed, we found significant correlations between within-category similarities in all three regions, in both the young and older adult groups.

Significant relationships between within-category similarities across the visual, auditory, and motor regions are consistent with the hypothesis that there may be a shared process or mechanism that leads to less reliable (noisier) activation patterns in older adults across the three domains. Aging is associated with several biological mechanisms that could plausibly interfere with the normal function of neurons throughout the brain, including free radical damage and oxidative stress (Harman, 1980), as well as damage to DNA and DNA repair mechanisms (Freitas & De Magalhães, 2011). Another possibility is age-related changes in neurotransmitter systems. Both dopamine (Bäckman, Nyberg, Lindenberger, Li, & Farde, 2006) and gamma-aminobutyric acid (GABA) (Chamberlain et al., 2021; Lalwani et al., 2019) systems have been reported to be affected by age and to be associated with neural distinctiveness.

### Limitations

An important limitation of this study is that it is cross-sectional. While we ascribe the differences that we observed to age-related effects, there is also the possibility of cohort effects (for example, group differences in childhood experiences, education, nutrition) that could potentially influence the results but that are unrelated to age per se. Future work could address this concern by utilizing longitudinal samples to investigate the trajectories of neural representations within individuals over time.

Additionally, this study only included younger and older adults and did not include any middle-aged participants. To our knowledge, Park et al., (2012) and Cassady et al., (2020) are the only studies addressing distinctiveness across the adult lifespan (but see also Chan et al., (2014) who investigated brain network segregation across the lifespan). Using data from the Dallas Lifespan Brain Study, Park and colleagues (2012) assessed neural distinctiveness in adults aged 20 to 89, concluding that neural dedifferentiation progresses linearly across the lifespan.

Similarly, Cassady et al, (2020) observed neural dedifferentiation across the lifespan in both motor and somatosensory systems.

Finally, like most neuroimaging studies of aging, the results reported here are correlational. We therefore cannot draw any causal inferences, but can only identity associations (e.g., between age and distinctiveness, between distinctiveness in different brain regions).

## Conclusion

In summary, we found evidence for age-related declines in neural distinctiveness in the visual, auditory, and motor cortices. These decreases appeared to be driven by both decreases in within-category similarities and increases in between-category similarities. We also found that cross-region relationships between neural distinctiveness were driven by within-category similarities, suggesting that age-related declines in the reliability of neural activity occur in tandem across the brain. Taken together, these findings support the idea that age-related dedifferentiation is influenced by changes in both the reliability and confusability of neural activity as we age and that changes in reliability in different brain regions are related.

